# Flowers as fungal extended phenotypes: nectar yeasts obfuscate among-plant differences in nectar sugar concentration

**DOI:** 10.1101/2025.02.04.636411

**Authors:** Carlos M. Herrera, Conchita Alonso

## Abstract

Nectar-dwelling yeasts modulate the ecology of interactions between flowers and pollinators. We evaluate here the hypothesis that floral nectar traits, which are a central element in most plant-pollinator relationships, can largely represent the extended fungal phenotypes of nectar-dwelling yeasts. The following specific question is addressed, Do local genotypes of the specialist nectar yeast *Metschnikowia reukaufii* have the ability to obfuscate intrinsic individual variation among *Helleborus foetidus* plants in nectar sugar concentration ? An array of paired plant-yeast genotypes mimicking a natural field situation was created in the laboratory by inoculating field-collected nectar from different plant individuals with distinct yeast genotypes following a factorial design. Chemical analyses of nectar sugars before and after exposure to yeasts were performed using ion-exchange high performance liquid chromatography. Plant individual, yeast genotype, and their interaction all had strong, significant effects on sucrose concentration of *H. foetidus* nectar. Yeasts abolished 79% of intrinsic variance among plants in nectar sucrose concentration and profoundly reshaped patterns of intrinsic among-plant variation. Our results support the hypothesis that among-plant variation in nectar sugar concentration found by pollinators in the field can sometimes reflect the extended phenotypes of nectar yeasts more closely than intrinsic differences among the plants themselves.

## Introduction

Phenotypic differences among conspecifics are a central determinant of the opportunity for selection in natural populations. In the case of plants, however, certain traits can sometimes depend more on the expression of genes belonging to intimately associated fungi than to those of the plants themselves, exemplifying instances of fungal “extended phenotypes” (Dawkins 1982; see, e.g., Dawkins 2004, Hunter 2009, 2018, Hawkes et al. 2021, for discussion and reviews). When fungal modifications of plant phenotypes involve floral features there can be fitness consequences for the individual plants via interference with pollinator behavior and plant reproduction (Batra and Batra 1985, Jennersten 1988, Jennersten and Kwak 1991, Roy 1993, 1996, Gange and Smith 2005, González-Mas et al. 2023). Fungi known so far to account for modifications of floral phenotypes mostly are systemic endophytes, generally parasites or pathogens bearing antagonistic relationships with their hosts. Nevertheless, recognition in recent years of the influence of nectar-dwelling yeasts on the interaction between plants and pollinators through their effects on floral nectar and the pollinators themselves (Herrera et al. 2008, 2013, Schaeffer et al. 2014, 2017, 2019, Vanette and Fukami 2016, 2018, Rering et al. 2018, Pozo et al. 2020, de Vega et al. 2022) motivates the hitherto unexplored hypothesis that intraspecific variation in nectar traits naturally occurring in the field might actually reflect the extended phenotypes of nectar-dwelling yeasts.

This paper reports results of an experimental test of this hypothesis for the interaction between plants of *Helleborus foetidus* (Ranunculaceae) and the nectar yeast *Metschnikowia reukaufii* (Metschnikowiaceae, Saccharomycetales). An array of paired plant-yeast genotypes mimicking a natural field situation was created in the laboratory by inoculating nectar from different plant individuals with cells of different yeast genotypes. Sugar analyses of nectar samples before and after a period of yeast population growth will be used to answer the following specific question: Do local genotypes of the yeast have the ability to obfuscate intrinsic differences among plants in nectar sugar concentration ? Our results revealed that the identity of the nectar yeast genotypes hosted by individual plants can become more important than intrinsic plant traits in shaping the variation in nectar traits encountered by foraging pollinators in natural plant populations, which in turn supports the hypothesis that pollinators might actually be interacting with sets of extended phenotypes of the nectar yeasts living in the flowers.

## Material and methods

### Study system and field sampling

This study focuses on the early-blooming perennial herb *Helleborus foetidus* and the dominant yeast in its floral nectar, *Metschnikowia reukaufii. Helleborus foetidus* inflorescences are produced in early winter and flowers are pollinated by bumble bees (Herrera et al. 2001). Flowers produce a sucrose-dominated nectar (Vesprini et al. 1999, Canto et al. 2011) that ordinarily harbors dense populations of *M. reukaufii*, the colonizing inocula of which are brought to flowers by foraging bumble bees (Brysch-Herzberg 2004, Canto et al. 2008). Metabolic activity of these yeasts induce substantial changes in composition and concentration of nectar sugars (Herrera et al. 2008, Canto et al. 2008, 2011).

Field sampling was conducted during February-March 2015 in a large population of *H. foetidus* in the Sierra de Cazorla, Jaén province, southeastern Spain. Floral nectar for the experiment was obtained from 20 widely spaced *H. foetidus* plants whose inflorescences had been previously bagged at the bud stage to exclude pollinators. For each individual plant, nectar from many flowers was gathered into a single bulk sample and stored at -20°C until used. Ten *M. reukaufii* strains from the same population were randomly chosen for the experiment from a large collection of local strains gathered from nectar of *H. foetidus* flowers exposed to pollinators.

### Experimental and analytical procedures

Prior to the experiment, nectar samples were filter-sterilized using a polyvinylidene difluoride 0.22 µm filter. Genetic distinctness of the experimental *M. reukaufii* strains was verified by fingerprinting them using polymorphic nuclear DNA microsatellite markers (Seino et al. 2013), and will be referred to hereafter as “genotypes”.

The experimental layout consisted of a two-way factorial design, with *M. reukaufii* genotype (*N* = 10 levels) and *H. foetidus* individual (*N* = 20 levels) as main factors, and two replicates per each genotype x individual combination (*N* = 381 experimental units in total; a few plants had insufficient nectar for replication). Each experimental unit consisted of a 0.2 mL PCR tube containing 15 *μ*L of virgin, sterile nectar plus 1 *μ*L of *M. reukaufii* cell suspension taken from a fresh culture on standard liquid medium. To assess differences among yeast genotypes in initial cell density per experimental unit, two replicated control tubes were also set per genotype at the time of inoculation, each consisting of 15 *μ*L of 10% lactophenol cotton blue fixing solution (LCB hereafter) plus 1 *μ*L of *M. reukaufii* cell suspension. After inoculation, all experimental tubes were held at 25°C without agitation. Aliquots for chemical analyses were taken from each experimental unit at 24 h and 48 h after inoculation. These short periods were chosen to impede extensive sugar depletion of nectar (Herrera et al. 2008), which would have biased comparisons.

Samples of virgin nectar from experimental plants (*N* = 40, two aliquots per plant) and all nectar samples taken in the course of the experiment (*N* = 762) were analyzed using ion-exchange high performance liquid chromatography. Analytical procedures and equipment were as described in Herrera et al. (2006) and Canto et al. (2011). Separate estimates of the concentration of the three sugars appearing in the samples (sucrose, fructose, glucose) were obtained by integrating areas under chromatogram peaks, and total sugar concentration was computed by summing partial figures. As previously found for *H. foetidus* nectar in the study area (Herrera et al. 2006, Canto et al. 2011), sucrose was virtually the only sugar in all nectar samples. It accounted on average for 99.2% (interquartile range = 99.4–100%) and 98.9% (interquartile range = 100–100%) of total sugars in virgin (pre-exposure to yeasts) and experimental (post-exposure) nectar samples, respectively. Only results for sucrose will be considered hereafter.

### Data analysis

Due to differences among source cultures, initial cell densities in experimental units differed among yeast genotypes (Chi-square = 28.7, df = 9, *P* = 0.0007, Kruskal-Wallis rank sum test; range of genotype means = 475–1399 cells/mm^3^), and experimental plants differed in the sucrose concentration of virgin nectar (*F*_19,20_ = 73.8, *P* = 1.7E-14; plant means range = 53– 77% weight-to-weight basis). To evaluate robustness of experimental results to this initial heterogeneity across experimental units, and to accurately dissect the role of yeast genotypes as agents of variation in nectar sugar across plants, two separate statistical analyses were carried out. In the first one, sucrose concentration in nectar of experimental units after exposure to yeasts was regressed on initial yeast cell density, initial sucrose concentration, and their interaction. In the second analysis, a linear model was fitted to post-exposure sucrose concentration using plant identity, yeast genotype and their interaction as predictors. Adding time of sample collection (24 h or 48 h) as a predictor had a negligible, statistically nonsignificant effect on results, and the two data sets were pooled for the analyses. In addition to estimating main effects and interactions in the two linear models, “relative importance” of each predictor was estimated by decomposition of the total variance (*R*^2^) accounted for by each model (Groemping 2007).

Statistical analyses were carried out using the R environment (R Core Team 2024). The raw data used here are available in Herrera (2025). Linear models were fitted using the lm function in stats package. Adequacy of fitted models was verified using the function check_model of the performance package (Lüdecke et al. 2021). Relative importance of predictors was obtained with function calc.relimp in package relaimpo (Groemping 2006). Marginal means and adjusted predictions (model-based estimates) of the response variable (sucrose concentration) were estimated from fitted models using the predict_response function in the ggeffects package (Lüdecke 2018). The average “blurring” effect of each yeast genotype on intrinsic among-plant variation in nectar sucrose concentration was inversely assessed with the correlation coefficient across plants between initial sucrose concentration in virgin nectar and the model-adjusted, predicted values of post-exposure concentration conditioned on the yeast genotype.

## Results

Exposure to yeasts largely wiped out initial interplant differences in sucrose concentrations. Initial heterogeneity across experimental units in yeast cell density and nectar sucrose concentration had negligible explanatory effects on variation in sucrose concentration after experimental nectar exposure to yeasts, as revealed by both the regression parameter estimates and the proportion of variance explained by single effects and their interaction (Table 1).

**Table 1.** Summary of statistical analyses testing for effects on nectar sucrose concentration after experimental exposure of nectar from 20 different plants to 10 yeast genotypes using a factorial design. Separate models were fitted to the data (*N* = 786 sugar concentration measurements) in which the predictors were initial yeast cell density and sucrose concentration, and plant and yeast genotype identities, respectively.

| Predictors in model | Effect | Parameter estimate (SE) | Test statistic | *P*-value | Variance explained (%) |
| --- | --- | --- | --- | --- | --- |
| Initial yeast and nectar features: |  |  | Student's *t* |  |  |
|  | Cell density (CD) | -0.0117 (0.009) | 1.35 | 0.18 | 2.1 |
|  | Sucrose concentration (SC) | 0.289 (0.13) | 2.26 | 0.024 | 9.1 |
|  | CD x SC | 0.0001 (0.001) | 0.84 | 0.39 | 0.084 |
| Plant and yeast identity: |  |  | *F* value (df) |  |  |
|  | Plant (P) |  | 31.6 (19, 586) | < 2E-16 | 32.5 |
|  | Yeast genotype (YG) |  | 31.7 (9, 586) | < 2E-16 | 15.3 |
|  | YG x P |  | 2.2 (171,586) | 7.9E-12 | 20.3 |

Plant and yeast identity, and their interaction, each had strong, statistically significant effects on nectar sucrose concentration after exposure to yeasts (Table 1). The proportion of variance explained by this model was remarkably high (*R*^2^ = 68.1%) given the large number of samples. Estimated contributions of main effects (plant and yeast identity) and their interaction to the variance explained by the model declined in the direction plant (32%) – plant x yeast (20%) – yeast (15%) (Table 1). Model-adjusted, predicted values of nectar sucrose concentration after exposure to yeasts varied among plants (range = 31–53%, conditioned on strain 1_A_10.2) and yeast strains (range = 40–62%, conditioned on plant CV01), but the strong plant x yeast interaction effect renders these figures only weakly informative (Figure 2).

The high explanatory value of the yeast x plant interaction deserves particular consideration in the context of the hypothesis examined here. First, the comparative effect of different yeast genotypes on nectar sucrose concentration was remarkably different for different individual plants, as illustrated by extensive criss-crossing of lines in the interaction plot of within-plant standardized values (Figure 1). Second, as a consequence of the strong interaction the set of paired combinations plant x yeast genotype created a heterogeneous two-dimensional space of post-exposure sucrose concentrations (Figure 2). And third, yeast genotypes differed widely in their capacity to blur individual variation in nectar sucrose concentration, as shown by the broad range across plants of the correlation coefficient between mean sucrose concentration before and after the experiment (*r* range = 0.142–0.736, mean = 0.454). The activity of yeast genotypes obliterated between 46–98% (mean = 79%) of intrinsic variance among plants in nectar sucrose concentration (estimated as 1 – *r*^2^).

**Figure 1.**
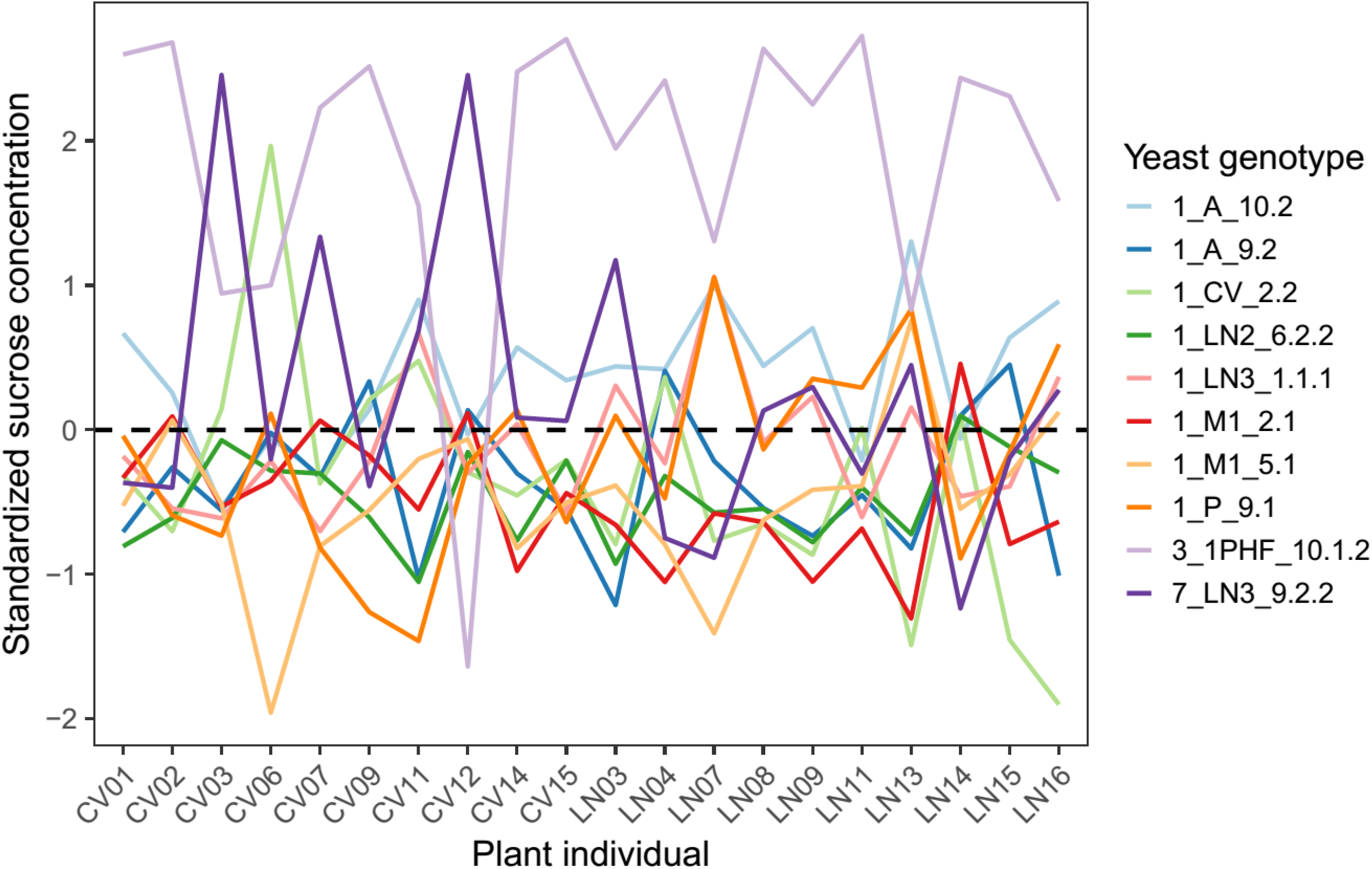
Interaction plot depicting differences in sucrose concentration after experimental exposure of nectar from different plants to different yeast genotypes. To emphasize the contrasting effect of the different yeast genotypes on different individual plants, and to facilitate comparisons across plants, model-predicted sucrose concentrations for each plant x yeast genotype combination were scaled and centered separately for each plant. See Figure 2 for the distribution of model-predicted values of sucrose concentration over the two-dimensional space defined by the 20 plants and 10 yeast genotypes tested in the experiment.

**Figure 2.**
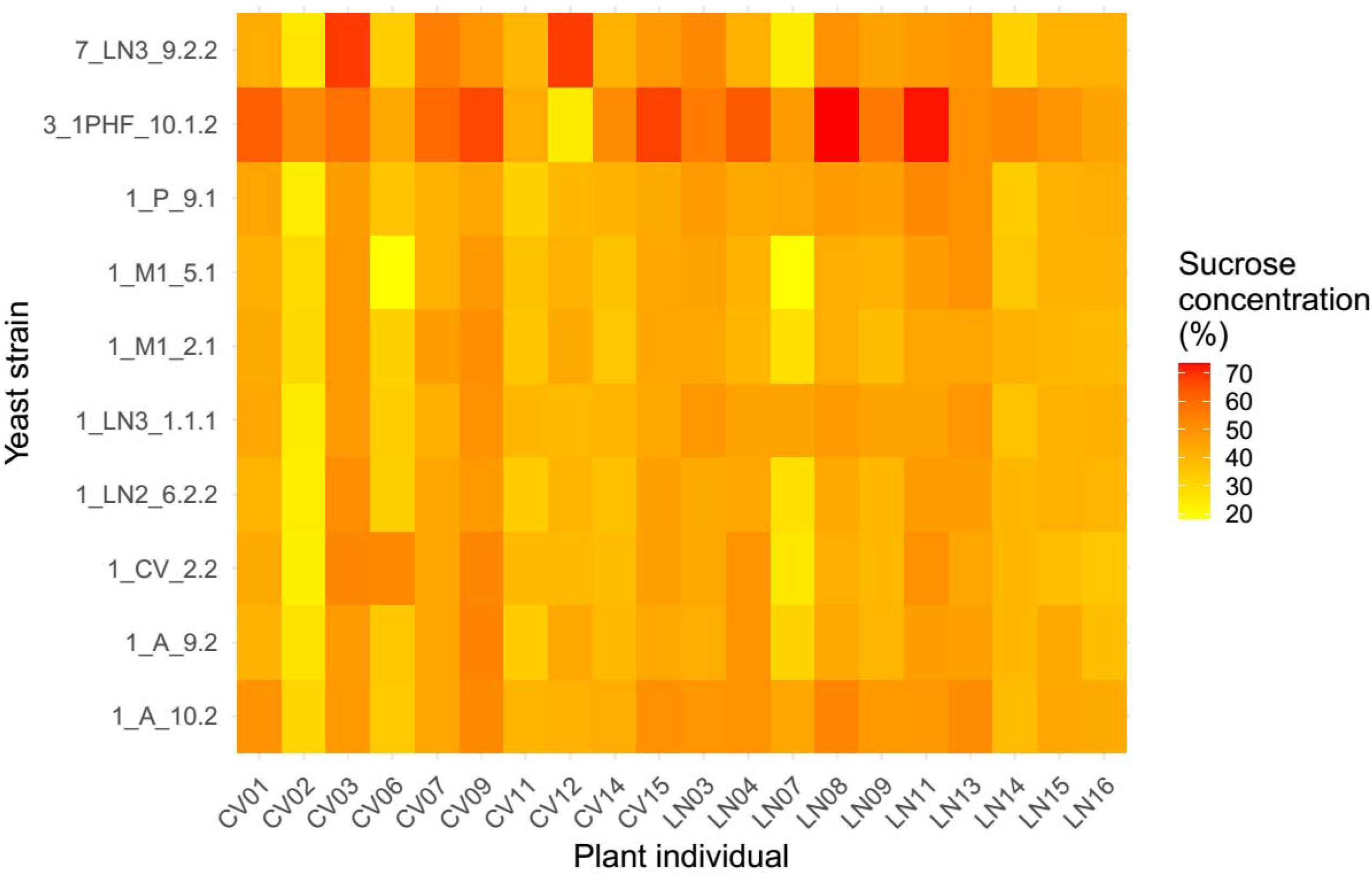
Model-adjusted, predicted sucrose concentration in nectar after exposure to yeasts for every plant identity x yeast strain combination.

## Discussion

Local *M. reukaufii* genotypes obfuscated intrinsic differences among individuals of *H. foetidus* in nectar sucrose concentration, a trait potentially important for the relationship with pollinators. After a relatively short exposure to the metabolic activity of *M. reukaufii* genotypes, among-plant differences in initial sucrose concentrations were nearly wiped out, while yeast identity and its interaction with plant identity acquired a major explanatory power. It must be stressed, however, that these experimental results probably underestimate the actual impact of yeasts on individual variation in sugar concentration in the field. On one side, initial yeast cell densities in experimental units were lower than the densities commonly found in nectar of *H. foetidus* flowers exposed to pollinators (Herrera et al. 2008, 2009). And on the other, length of exposure to yeasts in this study was shorter than the natural lifetime of individual *H. foetidus* flowers (up to 20 d; Herrera et al. 2001). Since sucrose concentration of *H. foetidus* nectar declines as yeast cell density rises with flower age (Herrera et al. 2008), yeast genotype effects in the field are probably larger than those reported here.

Applicability of the extended phenotype notion to nectar yeasts requires that the following three premises hold true: (1) yeast genotypes differ in physiological competence and ability to transform floral nectar, e.g., via changes in sugar composition or concentration; (2) in natural populations of a given plant, floral nectar of different individuals is inhabited by different yeast genotypes, so that plants might present to pollinators nectars with different features *because* they host different yeast genotypes; and (3) differences among coexisting plants in nectar traits of flowers exposed to natural conditions (i.e., visited by pollinators and colonized by yeasts) depend closely on the yeast genotypes living in their flowers. Premise 1 has been previously documented for *M. reukaufii* whenever metabolic differences among genotypes have been looked for (Herrera 2014, Pozo et al. 2015, Dhami et al. 2018), and premise 2 was supported by the finding that coexisting individuals of *H. foetidus* host distinct *M. reukaufii* genotypes in their floral nectar (Herrera et al. 2014). Premise 3 is supported by the results of the present study. Past and present results, taken together, thus provide compelling support for the hypothesis that natural variation in sugar concentration occurring in populations of *H. foetidus* can reflect the extended phenotypes of the nectar-dwelling yeast *M. reukaufii*. The potential implications of this finding for the interaction between *H. foetidus* and its pollinators are briefly discussed below.

Floral nectar is a major pollinator reward that strongly influences plant-pollinator interactions, and nectar features have been mostly interpreted in the light of selection from pollinators (Nicolson et al. 2007, Abrahamczyk et al. 2017). More recently, the potential role of nectar microorganisms in nectar evolution has begun to be acknowledged, although our understanding of their actual role is still in an incipient stage (Parachnowitsch et al. 2019, Barberis et al. 2024). Results of this study have revealed an hitherto unrecognized role of specialized nectar yeasts in nectar trait evolution: by reshaping intrinsic individual differences among plants, *M. reukaufii* yeasts will reduce the opportunity for pollinator selection on *H. foetidus* nectar sugar concentration. Such reduction will hinder selection on the plants’ intrinsic values of nectar sugar content, insofar as that variation is overshadowed by the transformative action of the yeast genotypes on nectar. Such reduction in opportunity for selection, however, will be in a sense compensated by new opportunities on an emergent axis. The variation among yeast genotypes in sugar transformative capacity, plus the strong plant x yeast genotype interaction effect on sugar concentration, will create opportunities for selection on sugar concentration over the two-dimensional space defined by combinations of plant x yeast genotypes (Figure 2). In this proposed scenario, pollinator selection on nectar traits of plant x yeast combinations will supersede “naive” pollinator selection on intrinsic nectar traits of individual plants, since the plant nectar phenotypes encountered by pollinators in the field will often be extended phenotypes of the yeast genotype(s) within the nectar. Nectar features could thus be considered as extended phenotypes of yeasts insofar as they are “candidate adaptations for the benefit of alleles responsible for variation in them” (Dawkins 2004), as discussed below.

Since initial conditions (sucrose concentration, yeast cell density) had a negligible effect on our experiment results, it seems safe to conclude that variation among experimental units in sucrose concentration after exposure to yeasts mainly arose from differences among yeast genotypes in growth rate and metabolic competence to exploit not just sucrose, but also other influential nectar constituents such as monosaccharides, proteins, amino acids and lipids which are also present in the nectar of *H. foetidus* but were not measured here (Herrera 1989, Vesprini et al. 1999). The concentration of these other nectar constituents most likely varied among plants, and in our experiment their collective, unmeasured effects are subsumed under the “plant identity” factor. Yeast genotypes with metabolic profiles variably suited to the chemical profile of a given plant’s nectar will probably grow and modify sucrose concentration at different rates (Herrera 2014). All else being equal, therefore, bumble bee discrimination for nectars with different sucrose concentrations (Cnaani et al. 2006, Whitney et al. 2008, Pozo et al. 2020) will translate into selection among plant-yeast combinations which have different sucrose concentrations (i.e., among different tiles in Figure 2). As bumble bees disperse yeast propagules and pollen grains simultaneously, any positive selection on a particular plant-yeast combination arising from pollinator discrimination will automatically benefit both the plant and the yeast genotype(s) involved, contribute to consolidate their association, increase its frequency, and perhaps eventually become “stabilized” within *H. foetidus* populations over the blooming season. In the long run, differential dispersal success of yeast genotypes associated with different plant individuals, and the likely stabilization of plant-yeast genotypic pairs postulated here, could eventually contribute to the genetic differentiation of *M. reukaufii* subpopulations found in flowers of *H. foetidus* individuals growing side by side, as well as the remarkable genetic and phenotypic diversity that characterize this yeast species (Herrera 2014, Herrera et al. 2014, Pozo et al. 2015, Dhami et al. 2018).

## Acknowledgments

We are grateful to Pilar Bazaga, Marina García and Esmeralda López-Perea for laboratory assistance; Ricardo Pérez for the development and implementation of sugar analytical procedures; and Consejería de Medio Ambiente, Junta de Andalucía por permission to work in the Sierra de Cazorla and providing invaluable facilities there. Different stages of this work were supported by grants CGL2010-15964 (Ministerio de Educación y Ciencia), P09-RNM-4517 (Junta de Andalucía), and PID2022-141530NB-C22 (Ministerio de Ciencia e Innovación).

## Data availability statement

Data, metadata and R script for analyses are available at figshare (https://figshare.com/s/2e27d89d49dc46e4adb1) (Herrera 2025)

